# Identification of novel mutations in RNA-dependent RNA polymerases of SARS-CoV-2 and their implications on its protein structure

**DOI:** 10.1101/2020.05.05.079939

**Authors:** Gyanendra Bahadur Chand, Atanu Banerjee, Gajendra Kumar Azad

## Abstract

The rapid development of SARS-CoV-2 mediated COVID-19 pandemic has been the cause of significant health concern, highlighting the immediate need for the effective antivirals. SARS-CoV-2 is an RNA virus that has an inherent high mutation rate. These mutations drive viral evolution and genome variability, thereby, facilitating viruses to have rapid antigenic shifting to evade host immunity and to develop drug resistance. Viral RNA-dependent RNA polymerases (RdRp) perform viral genome duplication and RNA synthesis. Therefore, we compared the available RdRp sequences of SARS-CoV-2 from Indian isolates and ‘Wuhan wet sea food market virus’ sequence to identify, if any, variation between them. We report seven mutations observed in Indian SARS-CoV-2 isolates and three unique mutations that showed changes in the secondary structure of the RdRp protein at region of mutation. We also studied molecular dynamics using normal mode analyses and found that these mutations alter the stability of RdRp protein. Therefore, we propose that RdRp mutations in Indian SARS-CoV-2 isolates might have functional consequences that can interfere with RdRp targeting pharmacological agents.

## INTRODUCTION

SARS-CoV-2 (a member of Coronaviruses) outbreak occurred in Wuhan, China in December 2019 and it became a pandemic by spreading to almost all countries worldwide. The SARS-CoV-2 causes COVID-19 disease that has created a global public health problem. As of May 8, 2020, more than 3.9 million confirmed COVID-19 cases were reported worldwide with 0.27 million confirmed deaths. To design antiviral therapeutics/ vaccines it is important to understand the genetic sequence, structure and function of the viral proteins. When a virus tries to adapt to a new host, in a new environment at a distinct geographical location, it makes necessary changes in its genetic make-up to bring slight modifications in the nature of its proteins(1). Such variations are known to helpviruses to exploit the host’s cellular machinery that promotes its survival and proliferation. The β lymphocytes of host’s adaptive immune system eventually identify the specific epitopes of the pathogenic antigen and start producing protective antibodies, which in turn results in agglutination and clearance of the pathogen(2–4). Being an efficient unique pathogen, a virus, often mutate its proteins in a manner that it can still infect the host cells, evading the host’s immune system. Even when fruitful strategies are discovered and engaged, the high rate of genetic change displayed by viruses frequently leads to drug resistance or vaccine escape(5).

The SARS-CoV-2 has a single stranded RNA genome of approximately 29.8Kb in length and accommodates 14 ORFs encoding 29 proteins that includes four structural proteins: Envelope (E), Membrane (M), Nucleocapsid (N) and Spike (S) proteins, 16 non-structural proteins (nsp) and 9 accessory proteins(6),(7) including the RNA dependent RNA polymerase (RdRp) (also named as nsp12). RdRp is comprised of multiple distinct domains that catalyze RNA-template dependent synthesis of phosphodiester bonds between ribonucleotides. The SARS-CoV-2 RdRp is the prime constituent of the replication/transcription machinery.The structure of the SARS-CoV-2 RdRp has recently been solved(8). This protein contains a “right hand” RdRp domain (residues S367-F920), and a nidovirus-unique N-terminal extension domain (residues D60-R249). The polymerase domain and N-terminal domain are connected by an interface domain (residues A250-R365).

For RNA viruses, the RNA-dependent RNA polymerase (RdRp) presents an ideal target because of its vital role in RNA synthesis and absence of host homolog. RdRp is therefore, a primary target for antiviral inhibitors such as Remdesivir(9) that is being considered as a potential drug for the treatment of COVID-19. Since, RNA viruses constantly evolve owing to the rapid rate of mutations in their genome, we decided to analyse RdRp protein sequence of SARS-CoV-2 from different geographical region to see if RdRp also mutate. Here, in the present study, we identified and characterised three mutations in RdRp protein isolated from India against that of the ‘Wuhan wet sea food market’(7) SARS-CoV-2. Altogether, our data strongly suggests the prevalence of mutations in the genome of SARS-CoV-2 needs to be considered to develop new approaches for targeting this virus.

## RESULTS

### Identification of mutations in RdRp protein present in Indian isolates

The SARS-COV-2 sequencing data was downloaded from NCBI (NCBI-Virus-SARS-CoV-2 data hub). There are 26 SARS-CoV-2 sequences available from India. We downloaded all Indian SARS-CoV-2 sequences and Wuhan SARS-CoV-2 sequence, which was deposited for the first time after COVID-19 cases started to appear in Wuhan province, China. We downloaded protein sequence of NSP12/ RdRp from the database. All these RdRp sequences were first aligned in CLUSTAL Omega to check for similarities or differences. The Clustal Omega algorithm produces a multiple sequence alignment by producing pairwise alignments. Using this algorithm, we identified seven mutations in isolates from Indian SARS-CoV-2 samples compared to Wuhan SARS-CoV-2 RdRp sequence as shown in figure 1. We considered the original Wuhan sequence as the wild type for this comparison. These mutations on RdRp are A97V, A185V, I201L, P323L, L329I, A466V and V880I respectively. Out of these, one isolate (QJQ28357) have triple mutation, two isolates (QJQ28429, QJQ28417) have double mutation, and rest have single mutations (figure 1).

**Figure 1:**
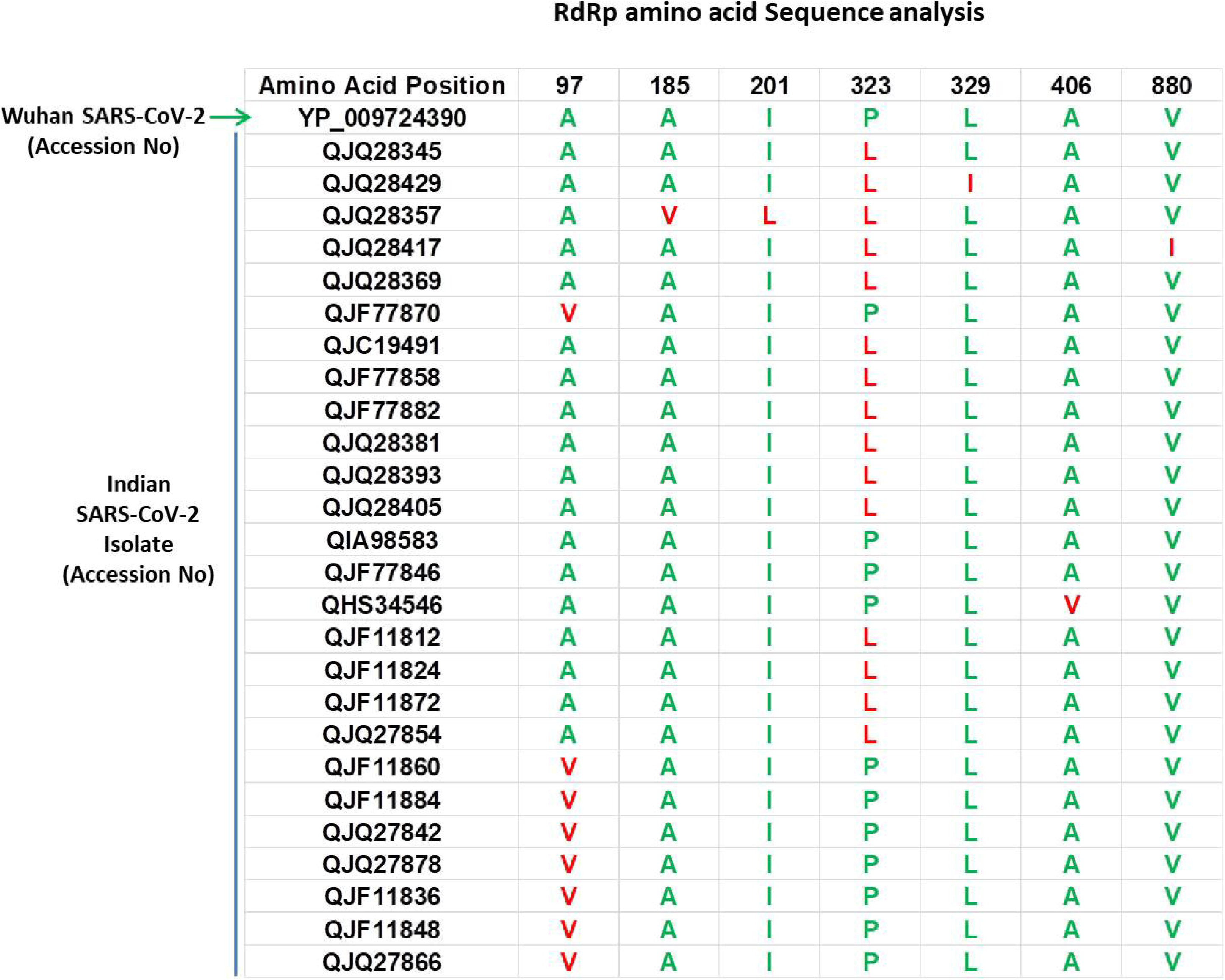
Multiple sequence alignment of Wuhan SARS-CoV-2 RdRp protein with sequences obtained from India. The mutations are highlighted in red font. Only those sequences are shown that have variations, rest of the sequences are identical among all samples.

### P323L mutation causes the alteration in secondary structure of RdRp

Next, we studied the effect of these mutations on secondary structure of RdRp. Our data revealed that mutation at four positions have no effect in secondary structure that includes I201L, L329I, A466V and V880I (figure 2 C, E, F and G). However, mutation in rest of the three sites causes change in secondary structure (A97V, A185V and P323L) as shown in figure 2 (panel A, B and D). At two positions (97 and 185) the alanine amino acid is substituted by valine. The valine side chain is larger than alanine, and substitution of valine at position 97 and 185 impairs packing of the protein as revealed by our secondary structure predications at these two sites (figure 2A and 2B, compare panel i and ii). Our analysis showed that there is addition of two sheet at positions 97 and 98 due to mutation of A97V (figure 2A, compare panel i and ii). Similarly, in A185V mutant, there is a loss of turn at 184 and replacement of helix with sheet structure at 181, 182, 183 and 184 positions (figure 2B, compare panel i and ii). Further, our secondary structure prediction also showed changes in secondary structure when proline is substituted by leucine at 323 position (Figure 2D and H, compare panel i and ii). The detailed analysis revealed that the mutant RdRp (P323L) have attained considerable changes in secondary structure at the mutation site. There is a loss of turn structures from position 323 and 324 and addition of five sheets at positions 321, 322, 323, 324 and 327 (Figure 2D and H, highlighted in green box). Proline possesses a unique property, its side chain cyclizes back on to the backbone amide position, due to which it contributes to the secondary structure formation because of its bulky pyrrolidine ring that places steric constraints on the conformation of the preceding residue in the helix. The substitution of proline to leucine in the mutant RdRp might result in loss of the structural integrity provided by proline. Leucine is a hydrophobic amino acid and generally buried in the folded protein structure. Altogether, the substitution of alanine to valine at position 97, 185 and proline to leucine at position 323 in mutant RdRp is changing the secondary structure of the protein that might have functional consequences.

**Figure 2:**
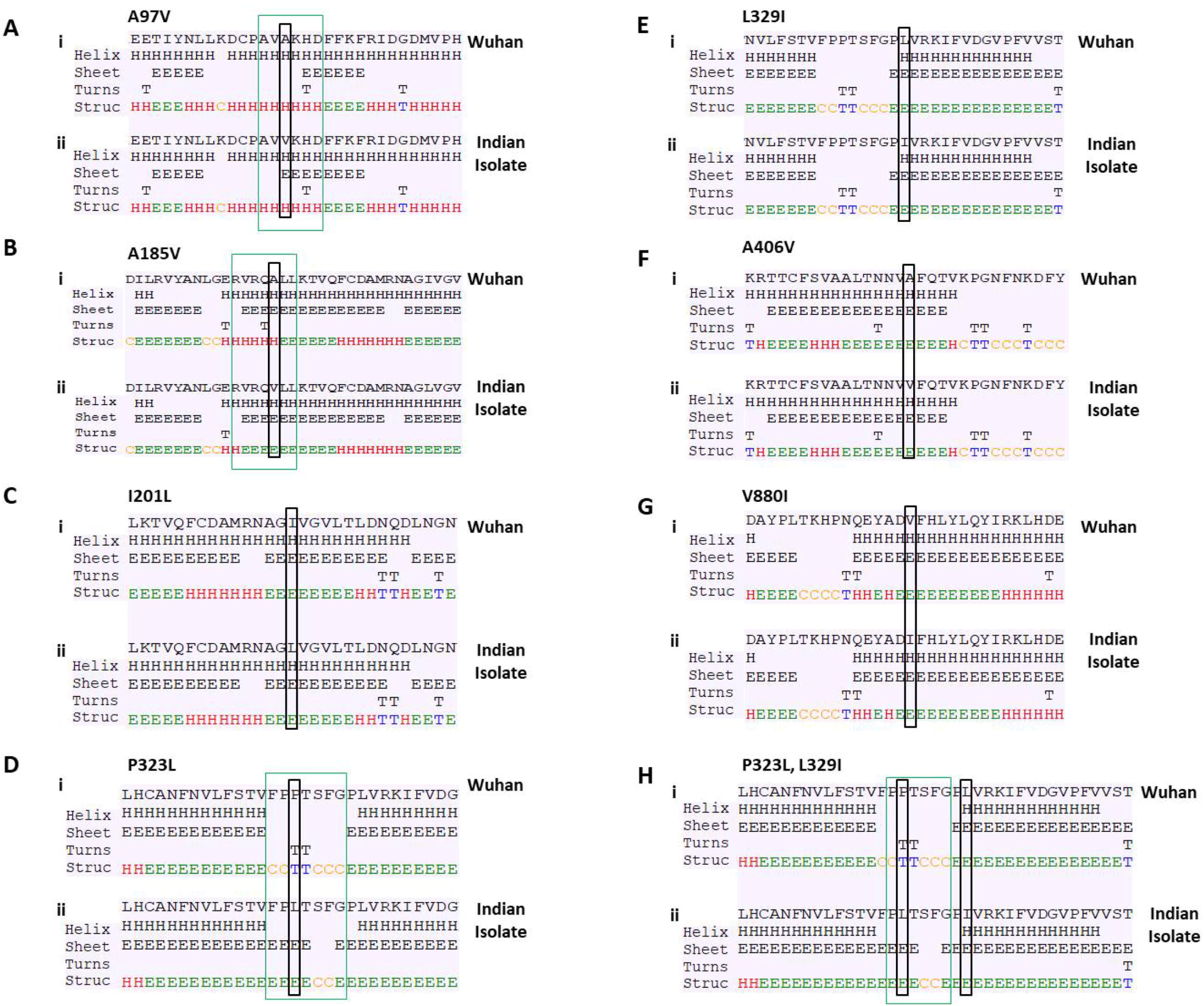
Effect of mutations on secondary structure of RdRp. Figure A, B, C, D, E, F, G and H demonstrate seven mutations observed in Indian isolates. Panel (i) represents sequence of Wuhan isolate and panel (ii) represents sequence of Indian isolates. The small rectangular box shows the mutant residue. The difference of secondary structure between Wuhan and Indian isolates are highlighted with green box.

### P323L alters the stability dynamics of tertiary structure of RdRp

To correlate the reflection of changes in secondary structure, if any, in the dynamics of the protein in its tertiary structure, we used DynaMut software(10) and performed normal mode analyses and studied protein stability and flexibility. Our data revealed that there is change in vibrational entropy energy (ΔΔS_Vib_ENCoM) between the wild type (Wuhan isolate) and the mutant Indian isolate (Figure 3A).Vibration entropy represents the major configurational-entropy contribution of proteins. By definition, it is an average of the configurational entropies of the protein within single minima of the energy landscape, weighted by their occupation probabilities(11). The negative ΔΔS_Vib_ENCoM of mutant RdRp represents the rigidification of the protein structure and positive ΔΔS_Vib_ENCoM represents gain in flexibility. Here, our data show that the mutation at A185V and P323L lead to rigidification of mutant protein structure Figure 3B and D). However, the mutation at I201L leads to increase in flexibility (Figure 3C). Further, we also calculated the free energy differences, ΔΔ*G*, between wild-type and mutant. The free energy differences, ΔΔ*G*, caused by mutation have been correlated with the structural changes, such as changes in packing density, cavity volume and accessible surface area and therefore, it measures effect of mutation on protein stability(12). In general, a ΔΔG below zero means that the mutation causes destabilization and above zero represents protein stabilization. Here, our analysis showed positive ΔΔ*G* for A185V and P323L mutations suggesting that P323L mutation is stabilising protein structure (Figure 3A); however, we observed negative ΔΔ*G* for I201L mutation indicating its destabilising behaviour (Figure 3A).

**Figure 3:**
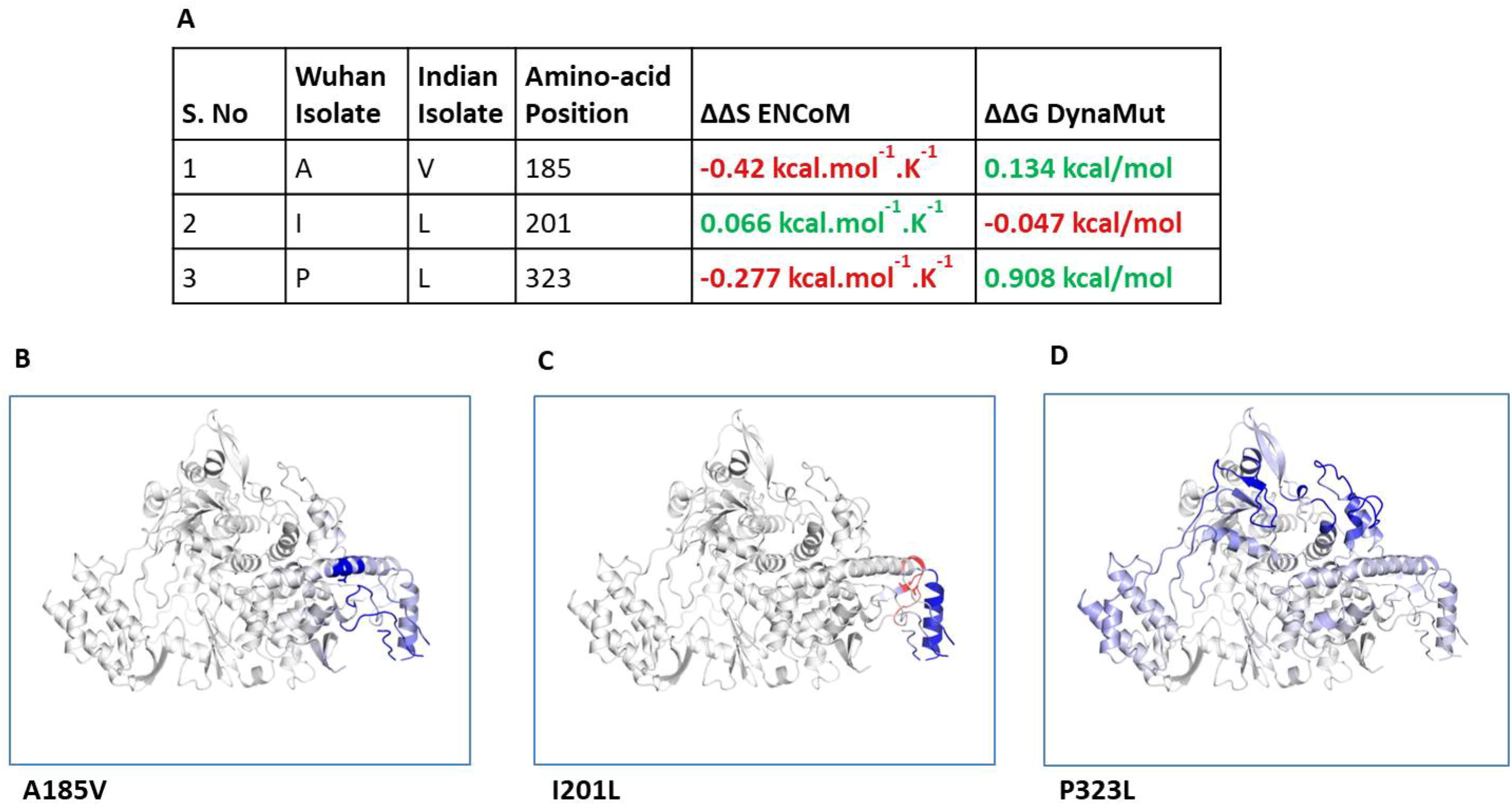
A) The table shows the values of change in ΔΔS ENCoM and ΔΔG due to the mutation. B) Δ Vibrational Entropy Energy between Wild-Type and Mutant RdRp, amino acids are colored according to the vibrational entropy change as a consequence of mutation of RdRp protein. **BLUE** represents a rigidification of the structure and **RED** a gain in flexibility.

We further closely analysed the changes in the intramolecular interactions due to these three mutations in RdRp. Our data showed that it is affecting the interactions of the residues which are present in the close vicinity of alanine, isoleucine and proline. The substitution of wild type residues with mutant residues alters the side chain leading to change of intramolecular bonds in the pocket, where these amino acid resides as shown in figure 4A, B and C. Therefore, it can be conclusively stated that the three mutations namely, A97V, A185V and P323L, in large, are changing the stability and intramolecular interactions in the protein that might have functional consequences.

**Figure 4.**
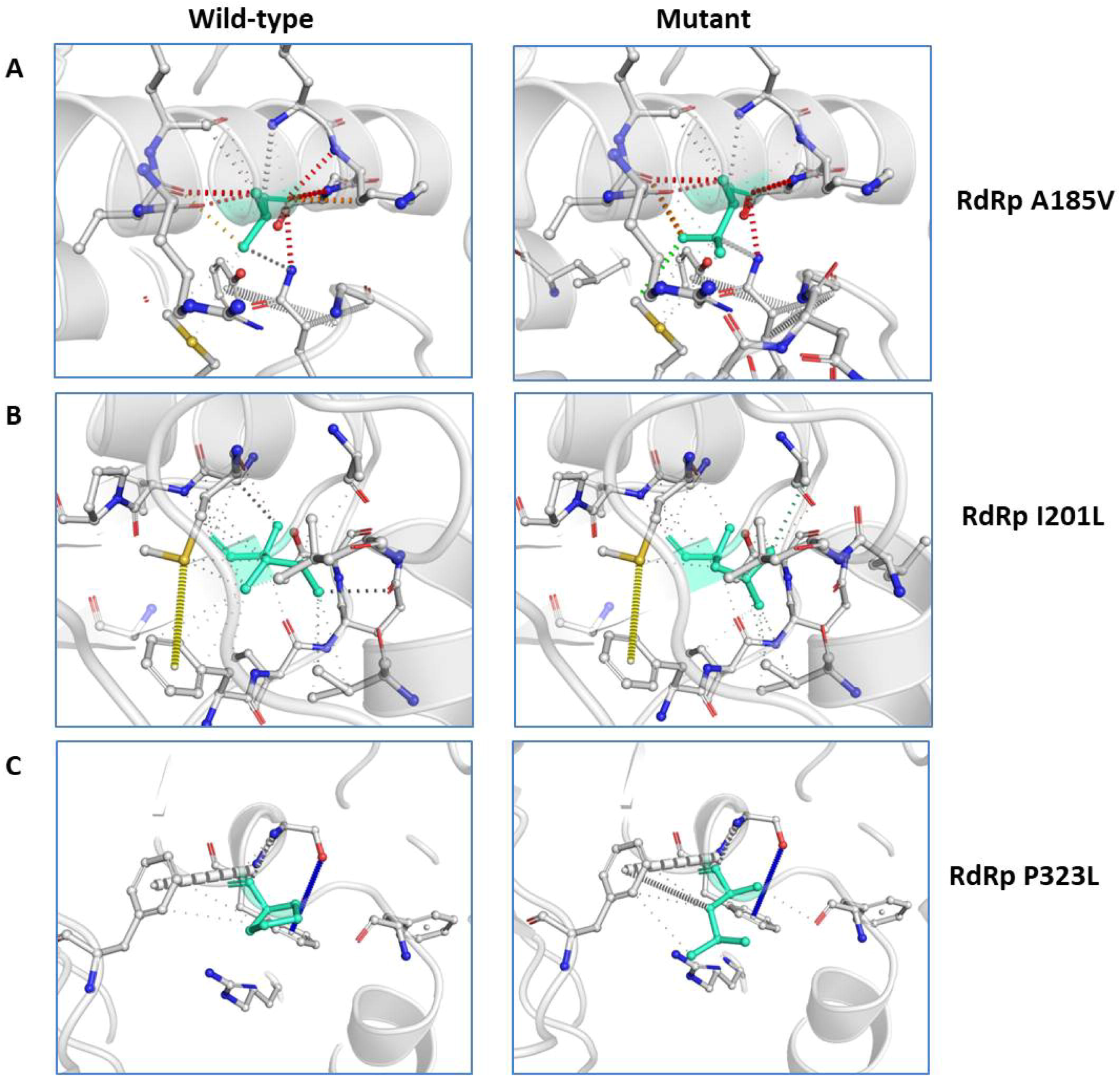
Interatomic interactions- (A, B and C) Wild-type and mutant residues are colored in light-green and are also represented as sticks alongside with the surrounding residues which are involved on any type of interactions.

## DISCUSSION

RNA viruses exploit all known mechanisms of genetic variation to ensure their survival. Distinctive features of RNA virus replication include high mutation rates, high yields, and short replication time (13). Mutation rates determine the amount of genetic variation generated in a population, which is the material upon which natural selection that can act (14). For this reason, a higher mutation rate correlates with a higher evolutionary rate, subject to natural selection. The mutation rates also determine the probability that a mutation conferring to drug resistance, antibody escape, or expanded host range may arise (15). Additionally, mutation rates can decide whether a virus population will become susceptible to drug-induced lethal mutagenesis.

The SARS-CoV-2 infected humans from Wuhan province in China and quickly spread to almost all countries worldwide. Since, this virus has already spread to different demographic areas having different climatic condition, temperature, humidity and seasonal variations; therefore, we can predict that this virus might be mutating to adapt to new environments. Towards this, we investigated the mutations in RdRp of SARS-CoV-2. We focused primarily on RdRp because it is an indispensable protein of this virus that helps in its replication and transcription. More importantly, there are many drugs which specifically target RdRp and are potent antivirals. Our study reports the RdRp mutations present in the Indian isolates of SARS-CoV-2 and, will help to understand the effect of variations on RdRp protein. Our data demonstrate that A97V, A185V and P323Lmutations lead to significant changes in the protein secondary structure (Figure 2). P323L mutation lies in the interface domain (residues A250-R365) of the RdRp protein. This domain helps in the coordination of N and C terminal domains of RdRp; therefore, a mutation in the interface domain might have drastic impact on the function of RdRp. A recent virtual molecular docking studies, screened approximately 7500 drugs to identify SARS-CoV-2 RdRp inhibitor revealed several potential compounds (DOI: 10.20944/preprints202003.0024.v1) namely, Simeprevir, Filibuvir and Tegobuvir, etc. Same study also predicted that these drugs bind RdRp at a putative docking site (a hydrophobic cleft) that includes phenylalanine at 326^th^ position.

Interestingly, the mutation identified in our study is very close to the docking site (P323L and L329I). Therefore, it is reasonable that the substitution of these two amino acids at 323 and 329 might interfere with the interaction of these drugs with RdRp. Further, our data also revealed that P323L mutation is causing stabilisation of the protein structure. The mutations in RdRp have already been linked to drug resistance in different viruses. Such as a study on RdRp of influenza A virus demonstrate that a K229R mutation confers resistance to favipiravir(16). Similarly, mutation in hepatitis C virus RdRp at P495, P496 or V499 & T389 and have been linked to drug resistance(17). Altogether, it is conceivable that many RNA viruses acquire drug resistance through mutations in RdRp. However, the functional characterisation of RdRp mutations investigated in our study needs to be carried out to understand the exact role of these mutations. These variations might lead to virus diversification and eventual emergence of variants or antibody escape mutants. Therefore, elaborate studies on SARS-CoV-2 protein sequence variations needs to be carried out that will help scientific community in better therapeutic targeting.

## METHODS

### Sequence retrieval

We downloaded all SARS-CoV2 sequences from the NCBI virus database as shown in table 1. As of now, there are 28 SARS-CoV-2 sequences from India have been deposited in this database, out of which, two sequences are not complete; therefore, we retrieved 26 datasets of SARS-CoV-2 (samples are from Indian). As a reference, we downloaded the sequence of SARS-CoV-2 that was first reported genome sequence deposited in the NCBI virus database from the ‘Wuhan wet sea food market area’ from the early days of COVID-19 pandemic(7). This virus was formerly called ‘Wuhan seafood market pneumonia virus having the accession number YP_009724390.

**Table 1:** Details of SARS-CoV-2 sequences used in the analysis

| S.No | Accession Number | Authors | Institute/ University |
| --- | --- | --- | --- |
| 1 | YP_009724390 | Wu,F., Zhao,S., et al., 2020 | Fudan University, Shanghai, China |
| 2 | QHS34546 | Yadav,P.D., et al., 2020 | NIV, Pashan, Pune, Maharashtra 411021, India |
| 3 | QIA98583 | Potdar,V., et al., 2020 | NIV, Pashan, Pune, Maharashtra 411021, India |
| 4 | QJC19491 | Pandit,R.,Shah,T., et al., 2020 | GBRC, Gandhinagar, Gujarat 382010, India |
| 5 | QJF11812 | Pattabiraman,C., et al., 2020 | NIMHANS, Bangalore, Karnataka 560029,India |
| 6 | QJF11824 | Pattabiraman,C., et al., 2020 | NIMHANS, Bangalore, Karnataka 560029,India |
| 7 | QJF11836 | Pattabiraman,C., et al., 2020 | NIMHANS, Bangalore, Karnataka 560029,India |
| 8 | QJF11848 | Pattabiraman,C., et al., 2020 | NIMHANS, Bangalore, Karnataka 560029,India |
| 9 | QJF11860 | Pattabiraman,C., et al., 2020 | NIMHANS, Bangalore, Karnataka 560029,India |
| 10 | QJF11872 | Pattabiraman,C., et al., 2020 | NIMHANS, Bangalore, Karnataka 560029,India |
| 11 | QJF11884 | Pattabiraman,C., et al., 2020 | NIMHANS, Bangalore, Karnataka 560029,India |
| 12 | QJF77846 | Muttineni,R., et al., 2020 | VRL, Dept. of Zoology, Osmania University, Hyderabad-500007,India |
| 13 | QJF77858 | Muttineni,R., et al., 2020 | VRL, Dept. of Zoology, Osmania University, Hyderabad-500007,India |
| 14 | QJF77870 | Muttineni,R., et al., 2020 | VRL, Dept. of Zoology,<br>Osmania University,<br>Hyderabad-500007,India |
| 15 | QJF77882 | Muttineni,R., et al., 2020 | VRL, Dept. of Zoology,<br>Osmania University,<br>Hyderabad-500007,India |
| 16 | QJQ27842 | Pattabiraman,C., et al., 2020 | NIMHANS, Bangalore,<br>Karnataka 560029,India |
| 17 | QJQ27854 | Pattabiraman,C., et al., 2020 | NIMHANS, Bangalore,<br>Karnataka 560029,India |
| 18 | QJQ27866 | Pattabiraman,C., et al., 2020 | NIMHANS, Bangalore,<br>Karnataka 560029,India |
| 19 | QJQ27878 | Pattabiraman,C., et al., 2020 | NIMHANS, Bangalore,<br>Karnataka 560029,India |
| 20 | QJQ28345 | Trivedi, P., et al., 2020 | GBRC, Gandhinagar,<br>Gujarat 382011, India |
| 21 | QJQ28357 | Hinsu, A., et al., 2020 | GBRC, Gandhinagar,<br>Gujarat 382011, India |
| 22 | QJQ28369 | Sabara, P., et al., 2020 | GBRC, Gandhinagar,<br>Gujarat 382011, India |
| 23 | QJQ28381 | Puvar, A., et al., 2020 | GBRC, Gandhinagar,<br>Gujarat 382011, India |
| 24 | QJQ28393 | Raval, J., et al., 2020 | GBRC, Gandhinagar,<br>Gujarat 382011, India |
| 25 | QJQ28405 | Gandhi, M., et al., 2020 | GBRC, Gandhinagar,<br>Gujarat 382011, India |
| 26 | QJQ28417 | Shah, T., et al., 2020 | GBRC, Gandhinagar,<br>Gujarat 382011, India |
| 27 | QJQ28429 | Pandya, M., et al., 2020 | GBRC, Gandhinagar,<br>Gujarat 382011, India |

### Sequence alignments and structure

All the RdRp protein sequences were aligned by multiple sequence alignment platform of CLUSTAL Omega(18). Clustal Omega is a multiple sequence alignment program that uses seeded guide trees and HMM profile-profile techniques to generate alignments between three or more sequences. The alignment file was carefully studied and differences in the amino acid changes were recorded.

### Secondary structure predictions

We used CFSSP(19) (Chou and Fasman secondary structure prediction), an online server, to predict secondary structures of SARS-CoV-2 RdRp protein. This server predicts the possibility of secondary structure such as alpha helix, beta sheet, and turns from the amino acid sequence. CFSSP uses Chou-Fasman algorithm, which is based on analyses of the relative frequencies of each amino acid in secondary structures of proteins solved with X-ray crystallography.

### RdRp dynamics study

To investigate the effect of mutation on the RdRp protein structural conformation, its molecular stability and flexibility, we used DynaMut software(10) (University of Melbourne, Australia). To run this software we first downloaded the known protein structure of SARS-CoV-2 RdRp from RCSB (PDB ID: 6M71(8)) and used it for analysis. Next, the 6M71 structure was uploaded on DynaMut software and effect of mutation in various protein structure stability parameters such as vibrational entropy; the atomic fluctuations and deformation energies were determined. DynaMut, a web server implements well established normal mode approaches that is used to analyse and visualise protein dynamics. This software samples conformations and measures the impact of mutations on protein dynamics and stability resulting from vibrational entropy changes and also predicts the impact of a mutation on protein stability.

## ACKNOWLEDGEMENTS

We would like to thank Patna University, Patna for providing necessary infrastructural support. No funding was used to carry out this research.

## CONFLICT OF INTEREST

Authors declare no conflict of interests.

## Notes

### Competing Interest Statement

The authors have declared no competing interest.

